# Clonogenic survival and relative biological effectiveness of pancreatic ductal adenocarcinoma under pristine and spread-out Bragg peak irradiation with protons and carbon ions

**DOI:** 10.64898/2026.09.30.754978

**Authors:** Janet M. Denbeigh, Alexandra Kneringer, Wilhelmina Radstake, Alessio Parisi, Sandra Barna, Shannon Hartzell, Sylvia Kerschbaum-Gruber, Ana Beatriz Dias, Patrick Fischer, Hermann Fuchs, Dea Slade, Dietmar Georg, Chris J. Beltran

## Abstract

Biological effectiveness in ion therapy varies with radiation quality and biological context, yet cancer cell response data acquired under well-characterized physical conditions remain limited. We investigated clonogenic survival and relative biological effectiveness (RBE) in PANC-1 and BxPC-3 pancreatic ductal adenocarcinoma (PDAC) cells irradiated at four positions along pristine 180.1 MeV proton and 352.4 MeV/u carbon ion Bragg curves, with 4 cm spread-out Bragg peak (SOBP) irradiation included as a clinically relevant comparison. Cell plane positioning and dose were verified in a water-tank system using an Advanced Markus ionization chamber, and Monte Carlo simulations using TOPAS provided dose-averaged linear energy transfer (LET_D_) and microdosimetric quantities. RBE was calculated relative to 200 kVp X-rays using the linear-quadratic survival model. Across the sampled pristine beam positions, spanning LET_D_ values of approximately 1-9 keV/µm for protons and 12-163 keV/µm for carbon ions, RBE increased with LET_D_. Proton RBE_10_ exceeded the conventional clinical value of 1.1 at distal positions, whereas carbon ion RBE_10_ increased from 2.3 to 5.2 in PANC-1 and from 1.3 to 3.7 in BxPC-3, with larger differences between cell lines observed near the Bragg peak. SOBP measurements showed different biological responses across field conditions despite comparable LET_D_. These physically characterized survival data provide PDAC-relevant benchmarks for evaluating whether RBE models reproduce biological responses across pristine and SOBP irradiation conditions, supporting the future integration of radiation-quality– dependent effects into ion planning.

## INTRODUCTION

Pancreatic ductal adenocarcinoma (PDAC) remains one of the most lethal malignancies worldwide, with five-year survival below 15% and limited curative options for patients with unresectable or locally advanced disease.^1^ Radiotherapy can contribute to local tumor control, but dose escalation is often constrained by the proximity of radiosensitive gastrointestinal organs. At the same time, intrinsic radioresistance contributes to tumor persistence and recurrence.^2^ Together, these anatomical and biological challenges have stimulated interest in ion therapy as a means of improving dose conformality while reducing exposure of surrounding normal tissues.^3-5^

Protons and carbon ions deposit dose with a characteristic depth-dose distribution that includes a Bragg peak and steep distal falloff, enabling greater sparing of normal tissues beyond the target volume.^6^ In addition to this physical advantage, carbon ions produce densely ionizing, high-linear energy transfer (LET) tracks that induce complex, spatially clustered DNA damage that is less readily repaired than damage caused by conventional low-LET photon irradiation.^7-11^ The biological effect of proton and carbon ion irradiation is commonly expressed as relative biological effectiveness (RBE), typically defined relative to a reference photon radiation quality. RBE depends on factors including ion species, LET, dose, and biological endpoint, and is further influenced by cellular characteristics such as radiosensitivity, cell cycle distribution, and DNA repair capacity.^12^

Clinically, biological effectiveness is represented using either fixed RBE values or model-based calculations. Proton therapy typically uses a constant RBE value of 1.1,^13, 14^ while carbon ion treatment planning relies on models such as the Local Effect Model (LEM I)^15^ and the modified Microdosimetric Kinetic Model (mMKM)^16^ to calculate RBE-weighted dose. These approaches provide a practical framework for incorporating biological effectiveness into treatment planning but simplify the complex factors that influence RBE. As a result, they may not fully capture differences in biological response among tumor types and cell models under comparable physical irradiation conditions. Experimental measurements are therefore needed to better characterize how biological effectiveness varies across radiation qualities and biological systems and to support interpretation of RBE-weighted dose.

Pristine Bragg curves provide a controlled experimental framework for examining how cellular response changes along the beam path. Cells can be irradiated at discrete depths spanning changes in dose-averaged LET (LET_D_) and microdosimetric characteristics, allowing biological response to be evaluated in relation to well-characterized physical conditions. However, physically characterized proton and carbon ion response data remain limited in PDAC, particularly across defined positions along pristine Bragg curves.^11, 17-19^ Complementary measurements under spread-out Bragg peak (SOBP) irradiation can extend these observations to mixed particle and energy spectra that more closely approximate clinical beam delivery.

We therefore measured clonogenic survival in two human PDAC cell lines, PANC-1 and BxPC-3, irradiated with protons and carbon ions at defined depths along pristine Bragg curves within a water-tank system at the MedAustron Ion Therapy Center. SOBP irradiation provided a comparison with modulated beam conditions more representative of clinical delivery. Cell plane positioning and delivered dose were verified dosimetrically,^20^ and Monte Carlo simulations were used to determine absorbed dose, LET_D_, and microdosimetric spectra at each position. Survival curves were fitted using the linear–quadratic (LQ) model, and RBE values were calculated relative to 200 kVp X-rays at surviving fractions of 50%, 10%, and 1%. Together, these measurements establish a PDAC-specific benchmark dataset relating clonogenic survival and RBE to physically characterized pristine and SOBP irradiation conditions and provide a foundation for evaluating and refining RBE models for biologically informed ion therapy planning.

## MATERIALS AND METHODS

### Radiation exposure

All irradiations were performed at the MedAustron Ion Therapy Center (Wiener Neustadt, Austria). Cells were irradiated at four depths along pristine proton and carbon ion Bragg curves. Additional irradiations were performed at the centers of proton and carbon ion spread-out Bragg peaks (SOBPs) with approximately uniform absorbed-dose profiles.

Pristine proton irradiations were delivered using a 180.1 MeV beam with a measured depth of maximum dose (d_max_) of 21.0 cm in water and a dose rate of 1.28 ± 0.02 Gy/min. Carbon ion irradiations used a 352.4 MeV/u beam with a measured d_max_ of 21.6 cm and a clinical fluence rate on the order of 10^10^cm^-2^s^-1^. For pristine proton irradiation, the ripple filter was removed to preserve the narrow Bragg peak; the ripple filter remained in place for pristine carbon ion irradiation. The Bragg peak widths at 80% of d_max_ were 6.10 mm for protons and 2.75 mm for carbon ions. Irradiations were delivered using scanned pencil beams with a 20 × 20 cm^2^ field and 2-mm spot spacing.

Before each irradiation campaign, simulations were used to identify four discrete depths along each pristine Bragg curve, as described in the Supplementary Materials. These positions spanned LET_D_ ranges of approximately 1–9 keV/µm for protons and 12–163 keV/µm for carbon ions. At each condition, five dose levels were delivered, with additional 0 Gy controls and sham-handled samples; dose levels were adapted to radiation quality and sampling position and are provided in Supplementary Figure 1.

For the SOBP irradiations, cells were positioned at the centers of 4 cm proton and carbon ion SOBPs. Plans were generated in RayStation version 7.99 (RaySearch Laboratories, Stockholm, Sweden) using a 17×9 cm^2^ field. Ripple filters were used for both ion species to produce a uniform SOBP, with energy layers spaced at 1 or 2 mm intervals. Cells were positioned at an equivalent water depth of 80 mm, corresponding to the center of each 40 mm SOBP, which nominally extended from approximately 60 to 100 mm. Proton energies ranged from 66.5 to 135.6 MeV, and carbon ion energies from 170 to 230 MeV/u. Additional details of the SOBP planning and delivery approach have been reported by Lohberger et al.^21^

Irradiation with 200 kVp X-rays at MedAustron served as the reference radiation quality for RBE calculations. Photon irradiations were performed using a YXLON Y.TU 320-D03 X-ray unit (YXLON GmbH, Hamburg, Germany) operated at 200 kVp and 20 mA with 3-mm Be, 3-mm Al, and 0.3-mm Cu filtration, resulting in a dose rate of 1.3 Gy/min. The irradiation setup and dosimetric commissioning have been described previously.^21^

### Dosimetry and depth verification

Comprehensive dosimetric verification was performed before all biological irradiations to ensure accurate positioning relative to the Bragg peak and reliable dose delivery at the cell plane. Depth-dose measurements were acquired using an Advanced Markus parallel-plate ionization chamber (Type 34045, PTW, Freiburg, Germany) mounted inside a 30 × 30 × 30 cm^3^ water phantom (Type 41023, PTW) using a customized plastic holder. The phantom entrance window was positioned at isocenter, 65 cm downstream of the beam nozzle. Chamber positioning along the beam axis was controlled with 0.1 mm resolution using the built-in phantom mechanical depth caliper.^20^ The water-equivalent thickness of the phantom entrance window (0.355 mm) and the effective point of measurement of the Advanced Markus chamber (P_eff_= 1.06 mm) were incorporated into all depth calculations. Because the chamber effective point of measurement was located 1.06 mm deeper than the cell monolayer, the dose at the cell plane was determined from the chamber measurement using Monte Carlo–derived depth-dose gradients. The final delivered dose at the cell plane was used for all survival analyses. Bragg peaks were first localized using coarse depth steps and subsequently refined using progressively smaller increments until the peak position was determined with sub-millimeter precision.

Treatment plans were generated in RayStation (RaySearch Laboratories, Stockholm, Sweden) using an initial prescription of 1 Gy at a 10 cm reference depth. For each biological sampling position, the initial plan was delivered and the dose measured using the Advanced Markus chamber. The monitor units (MU) were then rescaled based on the measured dose to generate a position-specific plan delivering a nominal dose of 1 Gy at the corresponding cell depth. Higher dose levels were produced by proportional scaling of the monitor units from the verified 1 Gy plan. The resulting cell plane dose was used for all survival analyses.

A locking bar indexed on the treatment couch ensured reproducible phantom positioning across measurement sessions. All chamber readings were corrected for temperature, pressure, and beam quality according to clinical dosimetry protocols. During biological irradiation, an Advanced Markus ionization chamber was positioned adjacent to the cell plane for real-time verification of dose delivery and spatial registration.

### Water-tank irradiation geometry

Cells were irradiated in chamber flasks positioned at defined depths within a previously characterized water-tank irradiation system.^20^ Horizontal scanned pencil beams entered through the flask base, enabling reproducible alignment of the cellular monolayer with predefined sampling positions. The irradiation setup is shown in Figure 1.

**Figure 1.**
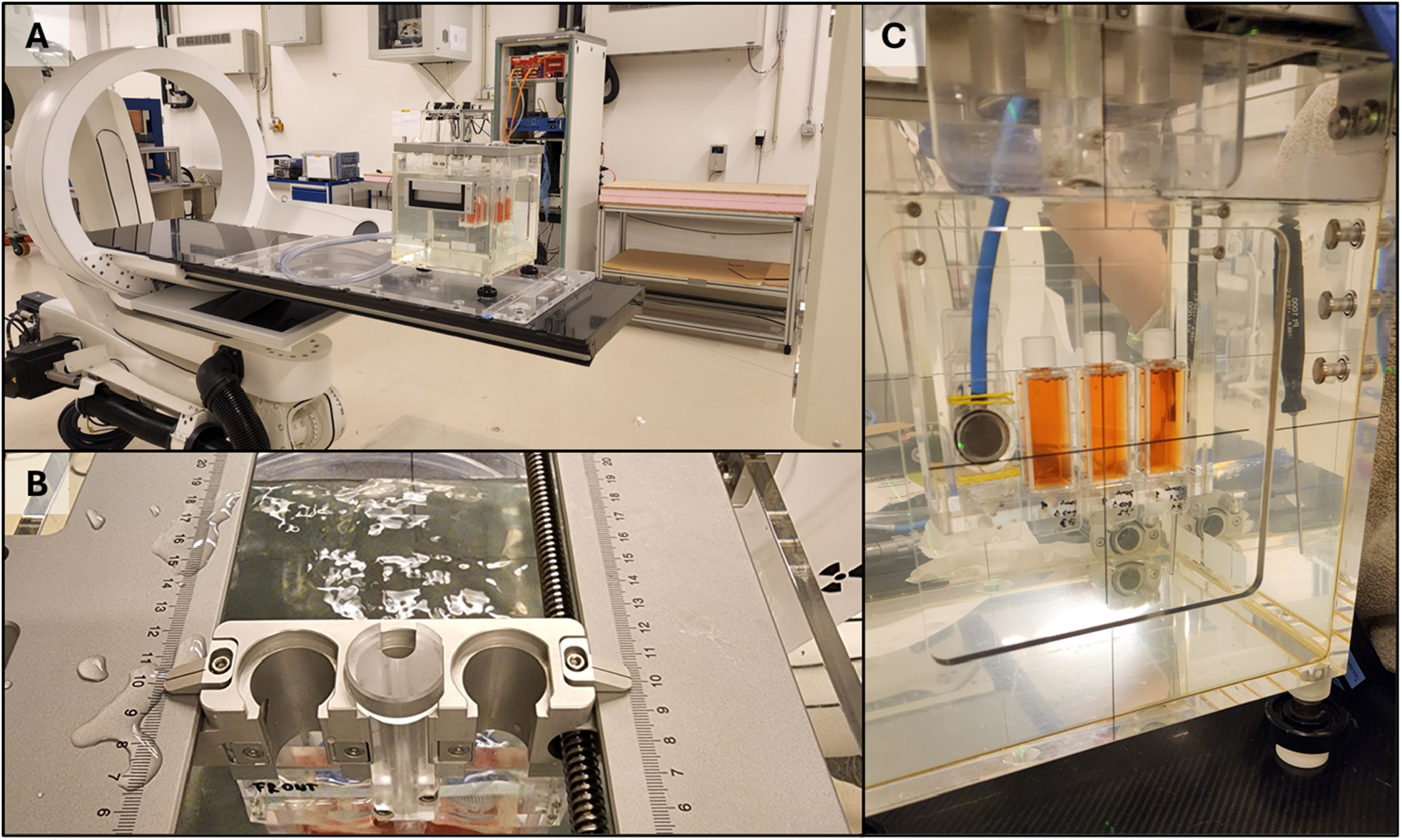
Water-tank irradiation setup for *in vitro* exposures and *in situ* dosimetric verification. (**A**) Water phantom positioned on the treatment couch and aligned with the horizontal beam axis. (**B**) Built-in mechanical depth caliper enabling longitudinal positioning with a 0.1 mm resolution (fine adjustment dial not shown). (**C**) Experimental configuration within the water tank showing three chamber flasks and an Advanced Markus parallel-plate ionization chamber placed adjacent to the cell plane for real-time verification of delivered dose during irradiation. **Alt text:** Three photographs show the water-tank irradiiaon setup, including the phantom aligned on the treatment couch, the mechanical depth-positioning system, and chamber flasks with an Advanced Markus ionization chamber positioned adjacent to the cell plane.

### Monte Carlo simulations and microdosimetry

Monte Carlo simulations were performed using OpenTOPAS v4.0^22^ to estimate absorbed dose, dose-averaged linear energy transfer (LET_D_), and the dose-mean lineal energy (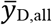) at each cell position (Figure 2 for pristine curves, Figure 3 for SOBP). Simulations were performed following completion of the irradiation campaigns using the experimentally determined beam and positioning parameters.

**Figure 2.**
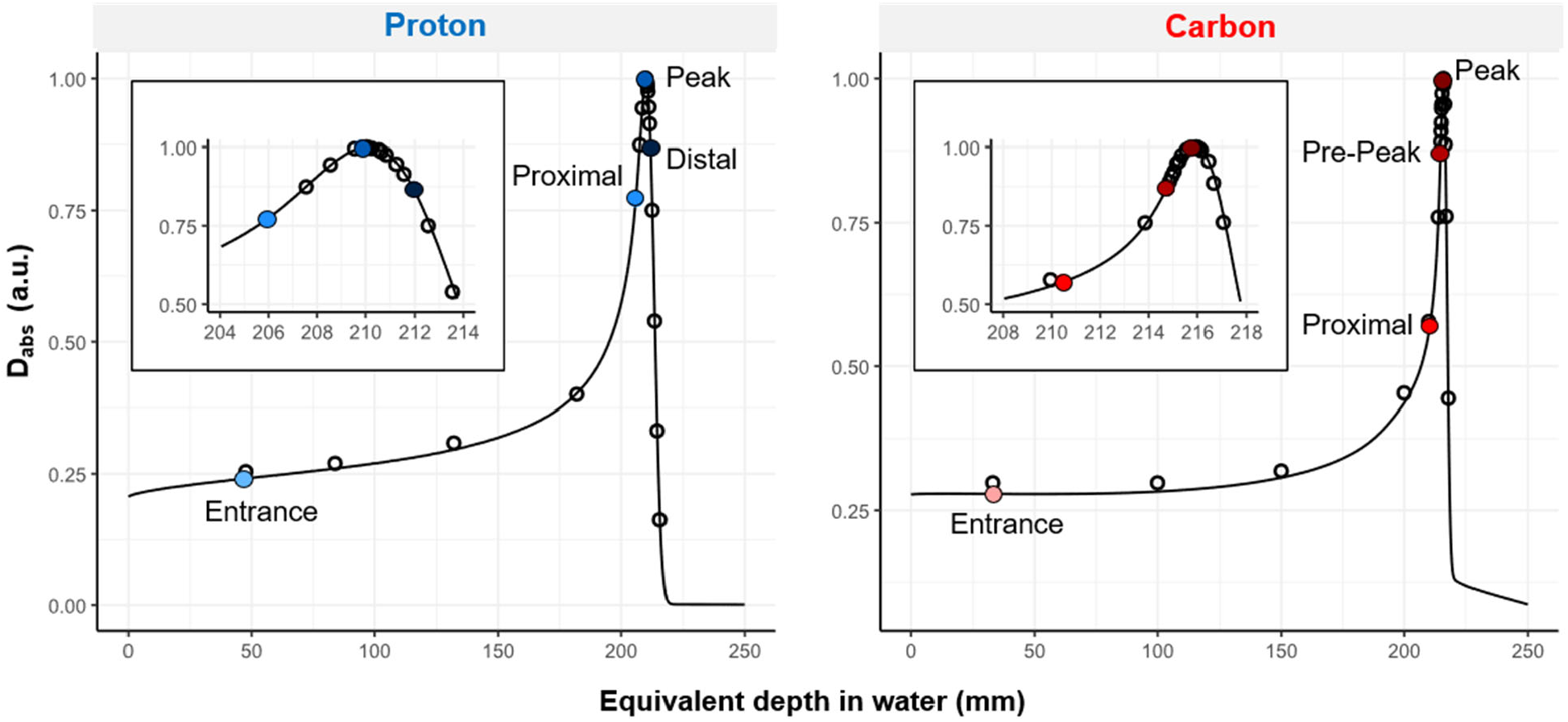
Normalized depth–dose profiles for monoenergetic proton and carbon beams with selected biological sampling positions. Pristine depth–dose distributions for the 180.1 MeV proton and 352.4 MeV/u carbon ion beam shown across the full range and magnified near the Bragg peak region (inset). Open circles represent measured depth– dose values acquired using the Advanced Markus ionization chamber; curves indicate Monte Carlo (TOPAS) simulations. Blue and red circles indicate the four cell exposure depths selected per beam. **Alt text:** Proton and carbon ion pristine depth–dose curves plotted against equivalent depth in water, with measured chamber data overlaid on Monte Carlo simulations. Four biological sampling positions are marked along each curve, spanning the entrance region through the Bragg peak or distal region. Insets enlarge the Bragg peak region.

**Figure 3.**
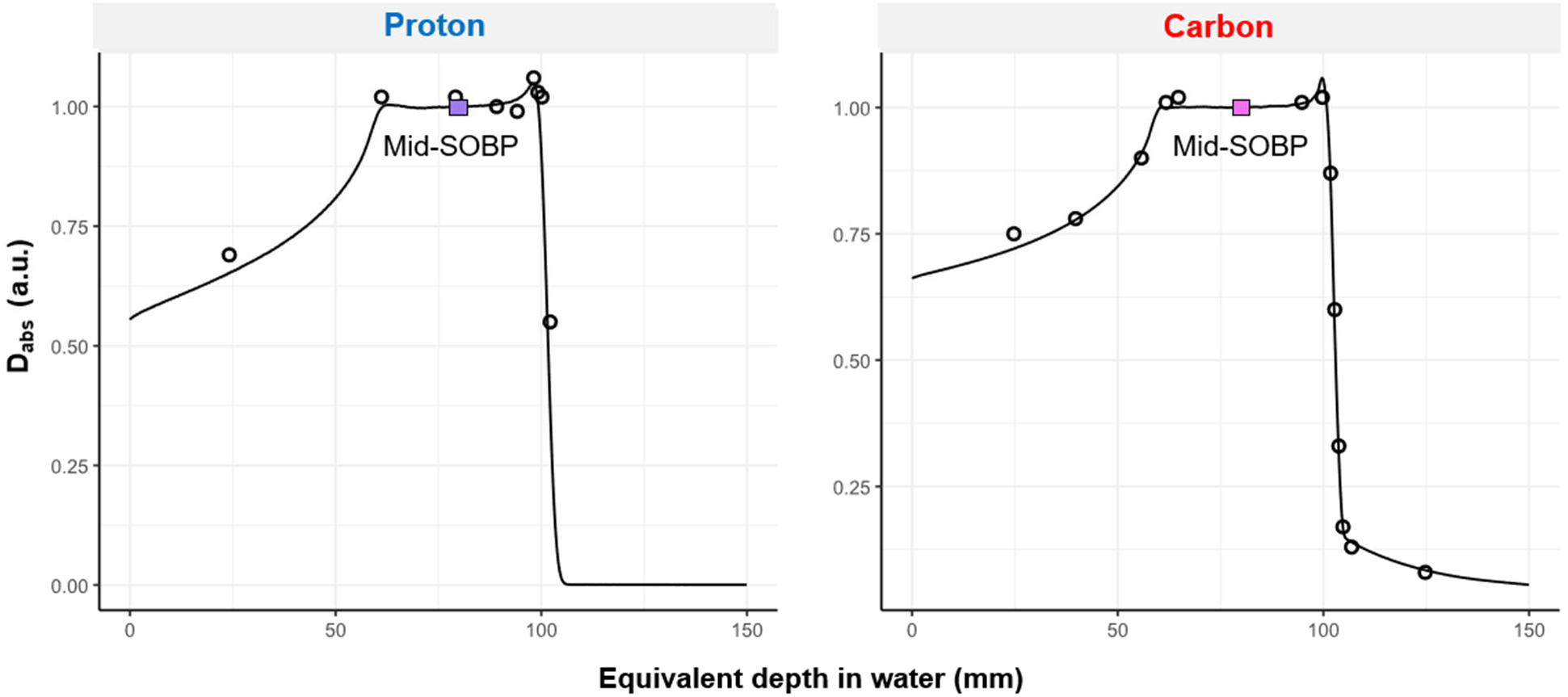
Normalized depth-dose profiles for the spread-out Bragg peak (SOBP) for proton and carbon ions with selected biological sampling positions. Open circles represent measured depth-dose values acquired using the Advanced Markus ionization chamber; curves indicate Monte Carlo (TOPAS) simulations. Purple or pink boxes indicate the cell positions during irradiation. **Alt text:** Proton and carbon ion spread-out Bragg peak depth–dose curves plotted against equivalent depth in water, with measured chamber data overlaid on Monte Carlo simulations. The biological sampling position is marked near the center of each approximately 40 mm wide SOBP.

Depth-dependent absorbed-dose profiles were calculated using the DoseToMedium scorer. A validated in-house scorer was used to calculate LET_D_ for four ion selections: (i) primary ions only; (ii) ions of the same type as the primary beam (^1^H or ^12^C ions); (iii) ions of the same element as the primary beam, including all hydrogen (H) or carbon (C) isotopes; and (iv) all ions. Unless otherwise stated, LET_D_ values used in the primary analysis were calculated for ^1^H ions for proton irradiation and ^12^C ions for carbon ion irradiation. LET_D_ values calculated using the alternative ion-selection criteria are reported in Table 1 and the Supplementary Materials.

**Table 1.** Physical beam characteristics and microdosimetric quantities at selected cell exposure positions. Equivalent depth in water is reported as mean ± positioning uncertainty for proton and carbon ion positions, including pristine and SOBP conditions. Dose-averaged linear energy transfer (LET_D_) is shown for primary ions only, ^1^H or ^12^C ions, all hydrogen or carbon isotopes, and all ions, together with dose-mean lineal energy (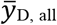). Values were derived from Monte Carlo simulations using experimentally validated beam parameters.

| Radiation type | Position | Depth in water<br>(mm) | $LET_{D,primary}$<br>(keV/μm) | $\overline{LET}_{D,1H \text{ or } 12C}$<br>(keV/μm) | $\overline{LET}_{D,H \text{ or } C}$<br>(keV/μm) | $\overline{LET}_{D,all}$<br>(keV/μm) | $\overline{y}_{D,all}$<br>(keV/μm) |
| --- | --- | --- | --- | --- | --- | --- | --- |
| **Photon**<br>200 kVp |  |  |  |  |  |  |  |
| **Proton**<br>180.1 MeV | Entrance | 47 ± 0.25 | 0.6 | 1.0 ± 0.1 | 1.0 ± 0.1 | 5.9 ± 0.2 | 7.7 ± 0.2 |
| | Proximal | 206 ± 0.25 | 3.0 ± 0.1 | 3.1 ± 0.1 | 3.1 ± 0.1 | 5.1 ± 0.1 | 7.6 ± 0.1 |
| | Peak | 210 ± 0.25 | 6.0 ± 0.3 | 6.1 ± 0.3 | 6.1 ± 0.3 | 7.0 ± 0.3 | 9.8 ± 0.3 |
| | Distal | 212 ± 0.25 | 8.4 ± 0.3 | 8.5 ± 0.3 | 8.5 ± 0.3 | 9.1 ± 0.3 | 11.9 ± 0.3 |
| 66.5-135.6 MeV | Mid-SOBP | 80 ± 0.25 | 2.9 ± 0.1 | 3.0 ± 0.1 | 3.0 ± 0.1 | 5.3 ± 0.2 | 7.6 ± 0.2 |
| **Carbon** |  |  |  |  |  |  |  |
| 352.4 MeV/u | Entrance | 33 ± 0.25 | 12.4 | 12.5 | 12.7 ± 0.1 | 12.8 ± 0.1 | 10.7 ± 0.1 |
| | Proximal | 210.5 ± 0.25 | 48 ± 1 | 49 ± 1 | 53 ± 1 | 46 ± 1 | 37 ± 1 |
| | Pre-Peak | 214.7 ± 0.25 | 105 ± 15 | 107 ± 15 | 109 ± 14 | 96 ± 14 | 86 ± 14 |
| | Peak | 215.8 ± 0.25 | 161 ± 19 | 163 ± 19 | 164 ± 18 | 145 ± 18 | 139 ± 18 |
| 170-230 MeV/u | Mid-SOBP | 80 ± 0.25 | 57 ± 2 | 57 ± 1 | 58 ± 1 | 53 ± 1 | 46 ± 1 |

Lineal energy distributions were computed using a revised implementation of an analytical microdosimetric function (AMF)^23, 24^ with a target radius of 0.3 µm. Additional details of the physical models, scoring methods, and computational implementation are provided in the Supplementary Materials.

### Cell culture

Human pancreatic ductal adenocarcinoma cell lines PANC-1 (ATCC CRL-1469) and BxPC-3 (ATCC CRL-1687) were maintained at 37 °C in a humidified incubator with 5% CO_2_. PANC-1 cells were cultured in Dulbecco’s Modified Eagle Medium (DMEM; Gibco 41965-062) and BxPC-3 cells in Roswell Park

Memorial Institute (RPMI)-1640 medium (Gibco 21875-091). Both media were supplemented with 10% fetal bovine serum (qualified, US origin; Thermo Fisher Scientific 26140-079; Lot. U2750588RP) and 1% penicillin–streptomycin (Sigma-Aldrich, P4333). Cells were passaged at 70–90% confluence using 0.05% trypsin-EDTA (Sigma-Aldrich, T3924) and used at passages below 10. Experiments were performed with asynchronously growing cell populations. Cultures were regularly tested and confirmed negative for mycoplasma contamination.

### Clonogenic survival assay

Twenty-four hours prior to irradiation, 4 × 10^5^ cells were seeded into chamber flasks (NUNC Lab-Tek 170920). Immediately before irradiation, the flasks were filled with prewarmed serum-free medium. Each irradiation condition was performed using three independently prepared flasks. Proton and carbon ion samples were positioned in the water tank as described above. The mean room temperature during irradiation was 21.9 ± 0.1 °C.

Irradiation times varied with beam condition and prescribed dose. The longest exposure was 14 min for the highest dose group at the first carbon ion position. For each position, sham-handled samples were placed in the water tank for a duration corresponding to the longest irradiation time used at that position. The mean irradiation time was 3 min 41 s (SD, 3 min 36 s). Experiments were completed across four separate irradiation sessions.

Immediately after irradiation, cells were trypsinized, counted, and reseeded into 6-well plates at dose- and cell-line-dependent densities, as detailed in Supplementary Table 1. Cells from each independently irradiated flask were distributed across six wells, yielding 18 wells per condition (six wells from each of three independently irradiated samples). Colonies were allowed to form for 13 days for PANC-1 and 16 days for BxPC-3.

Colonies were fixed with ice-cold 99.8% methanol (ACROS ORGANICS), stained with 0.5% crystal violet (ACROS Organics), gently rinsed with distilled water three times, and air-dried overnight. Colonies containing at least 50 cells were scored manually. Plating efficiency (PE) was defined as the ratio of colonies formed to seeded cells, and surviving fraction (S) was calculated by normalizing PE of each irradiated sample to the corresponding 0-Gy control.

Data from BxPC-3 cells at the proximal proton position were excluded because of insufficient plating efficiency in the corresponding unirradiated controls, which precluded reliable normalization and calculation of surviving fractions and RBE values.

### Statistical analysis and relative biological effectiveness calculations

All analyses were performed in R (Version 2025.09.1). Each irradiation condition included three independently prepared and irradiated flasks, with cells from each flask subsequently distributed across six wells for clonogenic assessment. Surviving fractions from the resulting 18 wells per dose condition were included in the survival-curve analysis. Surviving fractions (S) were linearized using the -ln(S) transformation. The linear-quadratic (LQ) model was fit using nonlinear least squares (nls, port algorithm) with non-negativity constraints on α and β. Fits were weighted by the inverse squared standard deviation of transformed S within each dose group. Error bars for surviving fractions represent propagated standard deviations accounting for variability in the treated samples and the average unirradiated control plating efficiencies used for normalization.

The estimated α and β parameters were used to calculate RBE at surviving fractions of 50%, 10%, and 1% (RBE_50_, RBE_10_, and RBE_1_, respectively) relative to the 200 kVp X-ray reference irradiation, using the following equation^25^:

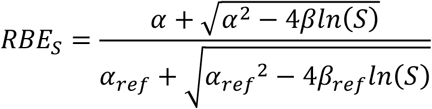

where S is the surviving fraction, α and β are the fitted LQ parameters for the ion irradiation condition, and α_ref_ and β_ref_ are the corresponding parameters for the reference X-ray irradiation, respectively. Standard errors for α and β were obtained from the nonlinear least-squares model fit and reflect uncertainty in the fitted parameter estimates. Uncertainties in the derived RBE values were estimated by propagation of these parameter standard errors.

## RESULTS

### Irradiation geometry and dosimetric verification

Monoenergetic proton and carbon ion beams produced the expected pristine Bragg depth–dose profiles (Figure 2). Chamber flasks were positioned at four predefined depths along each Bragg curve. Depth-dose mapping confirmed alignment of the selected cell positions with the intended regions of the Bragg curves. Concurrent Advanced Markus chamber measurements showed that the delivered cell plane dose was 99 ± 4% of the intended dose across all evaluated positions (mean ± SD).

Monte Carlo simulations reproduced the measured pristine depth–dose profiles (Figure 2) and provided estimates of LET_D_ and dose-mean lineal energy (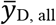) at each irradiation position (Table 1). The reported LET_D_ values varied according to the ions included in the averaging calculation, ranging from approximately 1–9 keV/µm for protons and 12–163 keV/µm for carbon ions. The measured SOBP depth–dose profiles were also reproduced by the Monte Carlo simulations, with cells positioned at the center of each 4 cm SOBP (Figure 3).

### Clonogenic growth characteristics

Both pancreatic cancer cell lines formed colonies suitable for quantitative clonogenic analysis (Figure 4). Although PANC-1 and BxPC-3 differed in colony morphology, assessment of plating efficiency across independently prepared samples showed generally consistent values within each experimental condition. The predefined colony-scoring criterion could be applied consistently to both cell lines. The exception was the BxPC-3 proximal proton condition, which was excluded because of insufficient plating efficiency in the corresponding unirradiated controls, as described above.

**Figure 4.**
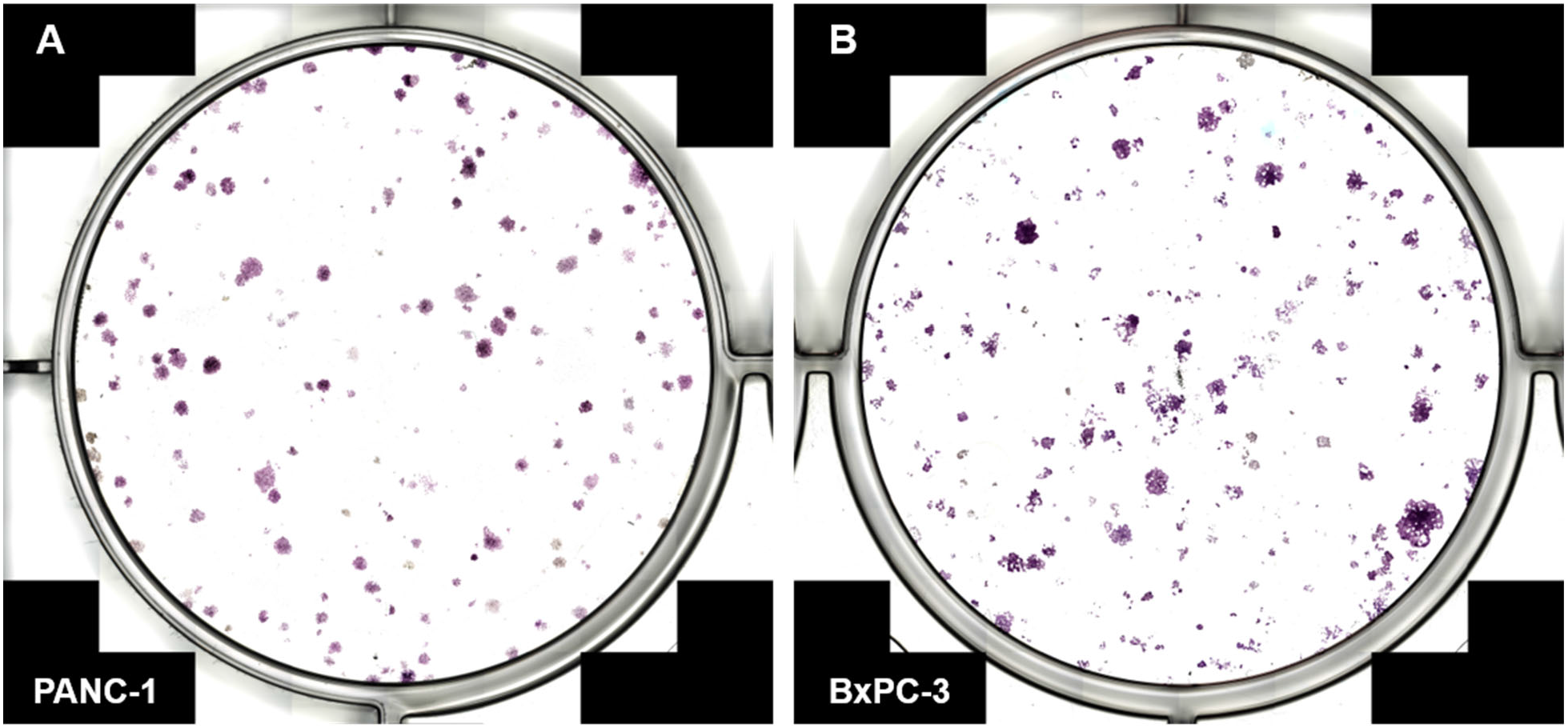
Representative colony morphology in control conditions. (**A**) PANC-1 colonies 13 days after irradiation (0 Gy control). (**B**) BxPC-3 colonies 16 days after irradiation (0 Gy control). Images were acquired at 2× magnification from individual wells of a 6-well plate following fixation and crystal violet staining, illustrating intrinsic differences in colony morphology between the two cell lines. **Alt text:** Representative crystal-violet-stained control wells show colony formation by PANC-1 and BxPC-3 cells. PANC-1 colonies are predominantly small and discrete, whereas BxPC-3 colonies show greater variation in size and morphology.

### Survival response as a function of dose-averaged linear energy transfer

Clonogenic survival varied with radiation quality and LET_D_ across the investigated conditions (Figure 5). Photon irradiation produced smoothly curving survival responses in both cell lines. For proton irradiation, survival was generally higher at the low-LET_D_ entrance positions and decreased toward the distal, higher-LET_D_ positions of the monoenergetic Bragg curve.

**Figure 5.**
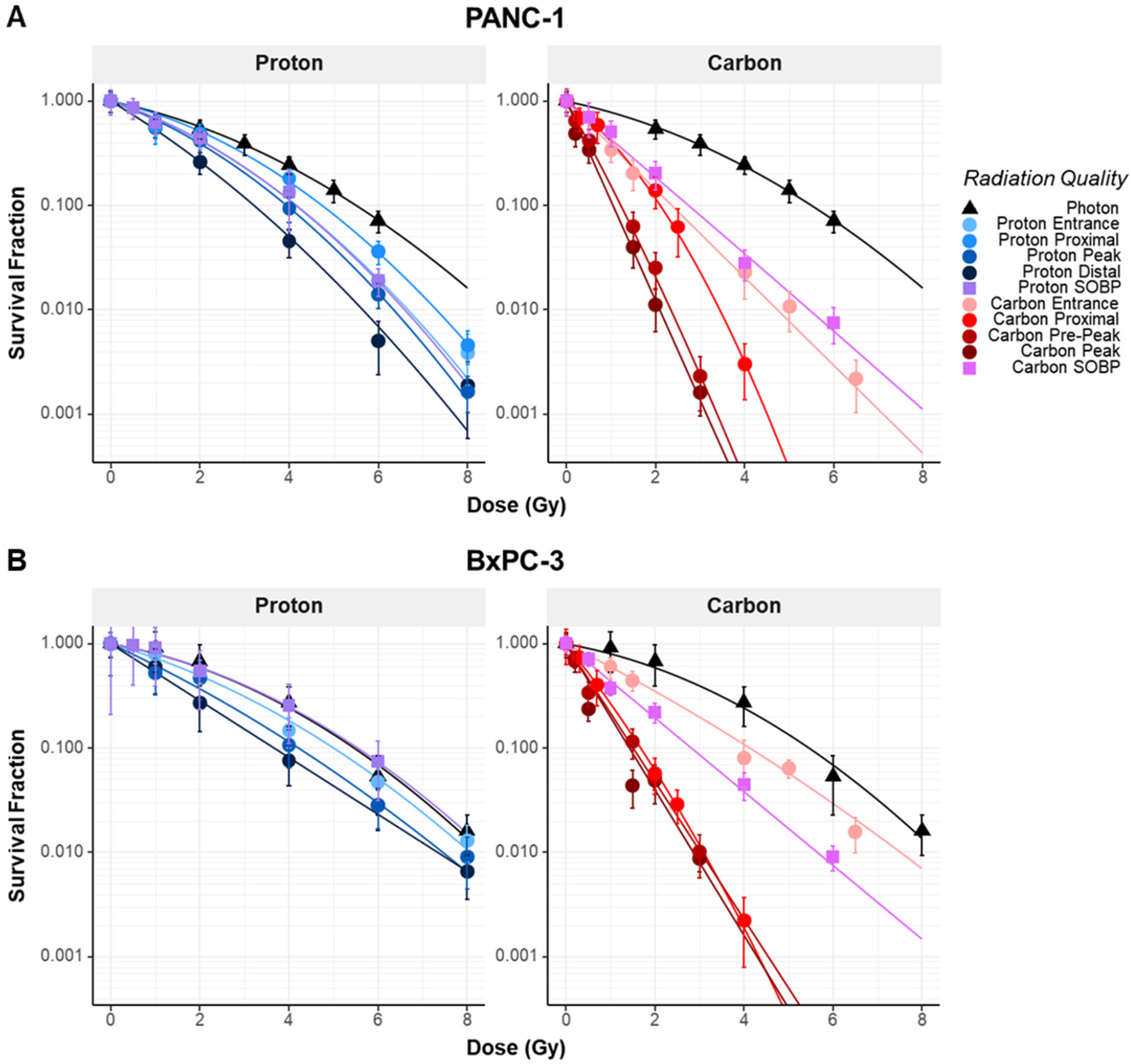
Clonogenic survival of PDAC cells as a function of absorbed dose and radiation quality. Survival curves for (**A**) PANC-1 cells and (**B**) BxPC-3 cells following irradiation with photons (black triangles), monoenergetic 180.1 MeV protons (blue circle shades), 352.4 MeV/u carbon ions (red circle shades), and proton and carbon SOBP (purple or pink squares, respectively). Symbols represent measured surviving fractions. Error bars represent propagated standard deviations, accounting for the variance in both the treated samples and the average unirradiated control plating efficiencies used for normalization. The y-axis is plotted on a logarithmic scale. Solid lines indicate linear–quadratic model fits. Progressive steepening of the survival curves is observed with increasing LET_D_ for both ion species, with the strongest effect at high-LET_D_ carbon peak positions. Data corresponding to the proximal proton sampling position for BxPC-3 cells were excluded from analysis due to insufficient plating efficiency in the associated unirradiated control samples, precluding reliable normalization of surviving fraction. **Alt text:** Dose–survival curves for PANC-1 (**A**) and BxPC-3 (**B**) cells following X-ray, pristine proton and carbon ion, and SOBP irradiation. Survival decreases with dose in both cell lines, with progressively steeper responses at higher-radiation-quality pristine positions and the steepest curves at high-LET_D_ carbon ion positions. The separation between carbon ion responses is greater in PANC-1 than in BxPC-3. In BxPC-3, the proton SOBP response closely follows the X-ray curve; the proximal proton condition is absent because it was excluded from analysis.

Carbon ion irradiation produced steeper survival curves than proton irradiation at comparable absorbed doses, with the lowest survival fractions observed at the higher-LET_D_ pre-peak and Bragg peak positions. The magnitude of this change, and the shape of the survival response differed between cell lines. At the higher-LET_D_ carbon ion positions, PANC-1 generally exhibited lower surviving fractions than BxPC-3 at comparable absorbed doses.

Survival following SOBP irradiation more closely resembled the entrance and proximal pristine beam responses than the high-LET_D_ pristine beam responses. This pattern was observed for both ion species and cell lines. Notably, in BxPC-3 cells, the proton SOBP survival curve closely overlapped the X-ray response.

### Linear–quadratic modeling and relative biological effectiveness estimation

Linear–quadratic fits described the observed survival trends across radiation conditions (Table 2). The fitted α and β parameters varied with radiation quality and LET_D_. At the highest LET_D_ carbon ion positions, α values were elevated and β approached zero, corresponding to survival curves with reduced curvature.

**Table 2.** Linear–quadratic model (LQ) parameter estimates and derived RBE values for PANC-1 and BxPC-3 cells. Fitted α and β parameters are reported as estimate ± SE. RBE values were calculated relative to 200kVp X-rays at surviving fractions of 50%, 10%, and 1% (RBE_50_, RBE_10_, and RBE_1_, respectively). SE represents the standard error of the fitted parameter estimate and reflects the precision of the model fit; it should not be interpreted as the standard deviation of experimental replicates or as measurement uncertainty of the raw survival data.

| | Radiation type | Position | $\alpha \pm \text{SE}$ | $\beta \pm \text{SE}$ | $\text{RBE}_{50}$ | $\text{RBE}_{10}$ | $\text{RBE}_1$ |
| --- | --- | --- | --- | --- | --- | --- | --- |
| **PANC-1** | **X-ray** | | $0.20 \pm 0.01$ | $0.039 \pm 0.002$ | | | |
| | **Proton** | Entrance | $0.32 \pm 0.02$ | $0.065 \pm 0.003$ | $1.4 \pm 0.1$ | $1.3 \pm 0.1$ | $1.3 \pm 0.0$ |
| | | Proximal | $0.22 \pm 0.01$ | $0.056 \pm 0.002$ | $1.2 \pm 0.0$ | $1.2 \pm 0.0$ | $1.2 \pm 0.0$ |
| | | Peak | $0.35 \pm 0.01$ | $0.059 \pm 0.003$ | $1.5 \pm 0.1$ | $1.4 \pm 0.0$ | $1.3 \pm 0.0$ |
| | | Distal | $0.59 \pm 0.02$ | $0.040 \pm 0.004$ | $2.2 \pm 0.1$ | $1.7 \pm 0.1$ | $1.5 \pm 0.1$ |
| | | Mid-SOBP | $0.31 \pm 0.02$ | $0.059 \pm 0.003$ | $1.4 \pm 0.2$ | $1.3 \pm 0.2$ | $1.3 \pm 0.3$ |
| | **Carbon** | Entrance | $0.97 \pm 0.03$ | $0.000 \pm 0.007$ | $3.3 \pm 0.1$ | $2.3 \pm 0.1$ | $1.8 \pm 0.1$ |
| | | Proximal | $0.70 \pm 0.06$ | $0.185 \pm 0.020$ | $2.9 \pm 0.2$ | $2.6 \pm 0.1$ | $2.5 \pm 0.1$ |
| | | Pre-Peak | $1.79 \pm 0.07$ | $0.075 \pm 0.031$ | $6.2 \pm 0.3$ | $4.5 \pm 0.2$ | $3.7 \pm 0.2$ |
| | | Peak | $2.20 \pm 0.07$ | $0.000 \pm 0.029$ | $7.5 \pm 0.3$ | $5.2 \pm 0.2$ | $4.1 \pm 0.2$ |
| | | Mid-SOBP | $0.83 \pm 0.03$ | $0.002 \pm 0.006$ | $2.8 \pm 0.1$ | $2.0 \pm 0.1$ | $1.6 \pm 0.1$ |
| **BxPC-3** | **X-ray** | | $0.17 \pm 0.03$ | $0.046 \pm 0.004$ | | | |
| | **Proton** | Entrance | $0.28 \pm 0.02$ | $0.036 \pm 0.003$ | $1.2 \pm 0.1$ | $1.1 \pm 0.1$ | $1.0 \pm 0.1$ |
| | | Peak | $0.45 \pm 0.02$ | $0.022 \pm 0.004$ | $1.7 \pm 0.1$ | $1.2 \pm 0.1$ | $1.1 \pm 0.1$ |
| | | Distal | $0.63 \pm 0.02$ | $0.000 \pm 0.004$ | $2.2 \pm 0.2$ | $1.4 \pm 0.1$ | $1.1 \pm 0.1$ |
| | | Mid-SOBP | $0.17 \pm 0.01$ | $0.044 \pm 0.003$ | $1.0 \pm 0.1$ | $1.0 \pm 0.1$ | $1.0 \pm 0.1$ |
| | **Carbon** | Entrance | $0.49 \pm 0.03$ | $0.017 \pm 0.006$ | $1.8 \pm 0.2$ | $1.3 \pm 0.1$ | $1.1 \pm 0.1$ |
| | | Proximal | $1.23 \pm 0.05$ | $0.086 \pm 0.018$ | $4.4 \pm 0.4$ | $3.2 \pm 0.2$ | $2.6 \pm 0.2$ |
| | | Pre-Peak | $1.52 \pm 0.07$ | $0.000 \pm 0.034$ | $5.2 \pm 0.5$ | $3.5 \pm 0.3$ | $2.7 \pm 0.2$ |
| | | Peak | $1.61 \pm 0.07$ | $0.000 \pm 0.026$ | $5.6 \pm 0.5$ | $3.7 \pm 0.3$ | $2.8 \pm 0.2$ |
| | | Mid-SOBP | $0.82 \pm 0.03$ | $0.000 \pm 0.006$ | $2.8 \pm 0.2$ | $1.9 \pm 0.1$ | $1.4 \pm 0.1$ |

Across the sampled pristine beam positions, RBE increased with increasing LET_D_. Proton RBE_10_ exceeded the conventional clinical value of 1.1 at the distal position, reaching 1.7 in PANC-1 and 1.4 in BxPC-3, whereas carbon ion RBE_10_ increased from 2.3 to 5.2 in PANC-1 and from 1.3 to 3.7 in BxPC-3 near the Bragg peak. Although both pancreatic cancer cell lines demonstrated LET-dependent biological enhancement, RBE was generally higher in PANC-1 than in BxPC-3. At the highest LET_D_ positions, β estimates approached zero.

For the SOBP conditions, proton RBE differed between the two cell lines. In PANC-1, proton SOBP values were similar to those measured at the pristine entrance position, with RBE_50,_ RBE_10_, and RBE_1_ values of 1.4, 1.3, and 1.3, respectively. In BxPC-3, the corresponding proton SOBP RBE values were approximately 1.0 at all three surviving fraction endpoints. Carbon ion SOBP irradiation produced RBE_50,_ RBE_10_, and RBE_1_ values of 2.8, 2.0, and 1.6 in PANC-1 and 2.8, 1.9, and 1.4 in BxPC-3.

### Relationship between relative biological effectiveness and dose-averaged linear energy transfer

Plotting RBE as a function of LET_D_ demonstrated a gradual increase across the sampled pristine proton positions and a steeper increase across the pristine carbon ion positions (Figure 6 for LET_D, 1H or 12C_, and Supplementary Materials for alternative LET_D_ definitions and 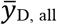). The magnitude of the RBE increase differed between PANC-1 and BxPC-3, particularly at the higher LET_D_ carbon ion positions.

**Figure 6.**
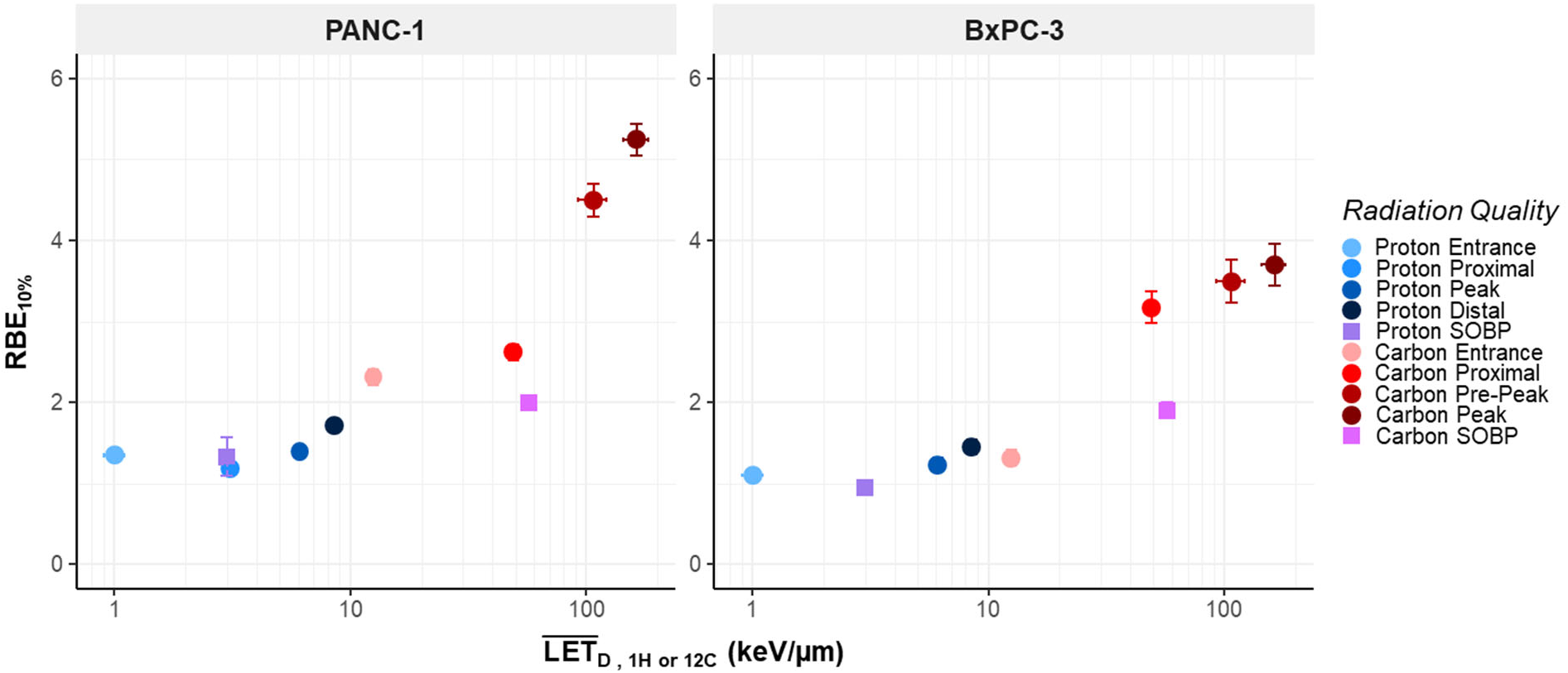
RBE_10_ as a function of LET_D,1H or 12C_for PANC-1 and BxPC-3 cells. Relative biological effectiveness at 10% surviving fraction (RBE_10_) plotted against Monte Carlo–derived dose-averaged LET for ions of the same type as the primary beam (^1^H or ^12^C ions). Blue symbols represent 180.1 MeV proton sampling positions (entrance, proximal, peak, distal), and red symbols represent 352.4 MeV/u carbon ion sampling positions (entrance, proximal, pre-peak, peak). Purple and pink squares represent SOBP proton and carbon results, respectively. Error bars reflect propagated uncertainty from the fitted LQ parameter estimates used in the RBE calculation. Across the sampled pristine beam positions, RBE_10_ increases with LET_D_ for both ion species, with a steeper relationship observed for carbon ions at higher LET values. **Alt text:** Scatter plots show RBE_10_ versus dose-averaged LETD for PANC-1 and BxPC-3 cells. RBE_10_ increases gradually across pristine proton positions and more steeply across pristine carbon ion positions, with higher values in PANC-1 at the highest carbon ion LETD. Proton and carbon ion SOBP measurements are shown alongside the pristine-beam data.

The SOBP measurements did not follow the pristine beam relationships. In PANC-1, proton SOBP RBE values aligned with the pristine proton entrance position, whereas the BxPC-3 proton SOBP values remained near 1.0. The carbon ion SOBP had an LET_D_ of approximately 57 keV/µm but produced lower RBE values than the proximal pristine carbon ion position, which had a similar LET_D_. Thus, similar dose-averaged LET_D_ values were not necessarily associated with similar RBE across pristine and SOBP irradiation conditions.

## DISCUSSION

### Biological response under pristine and spread-out Bragg peak irradiation

This study provides physically characterized clonogenic survival and RBE data for two PDAC cell lines irradiated at defined positions along monoenergetic pristine proton and carbon ion Bragg curves. Water-tank positioning, cell plane dosimetry, and Monte Carlo–derived LET_D_ and microdosimetric quantities allowed biological response to be evaluated across distinct regions of each pristine beam. Measurements at the centers of proton and carbon ion SOBPs provided a comparison with irradiation conditions more representative of clinical beam delivery.

Taken together, the measurements show that the relationship between radiation quality and biological effectiveness depends on both the irradiation field and the biological system. RBE increased across the sampled pristine beam positions, particularly for carbon ions, but comparable LET_D_ did not necessarily correspond to equivalent biological effectiveness under pristine and SOBP conditions. The differing responses between PANC-1 and BxPC-3 further indicate that radiation quality alone does not account for the observed variation in RBE.

### Ion-and field-specific differences in relative biological effectiveness

For proton irradiation, RBE_10_ remained close to the clinically assumed value of 1.1 across most pristine beam positions but increased at the distal edge, reaching 1.7 in PANC-1 and 1.4 in BxPC-3. This finding is consistent with previous experimental and modeling studies demonstrating increased proton biological effectiveness with increasing radiation quality toward the distal region of the beam.^14, 26^ Similar trends were observed at other survival fraction endpoints, with distal RBE_50_ values approaching 2.2 in both cell lines. The proton SOBP response differed between the cell line models: in PANC-1, the SOBP RBE_10_ was 1.3 and comparable to the pristine entrance region, whereas in BxPC-3 it remained near 1.0 and closely approximated the X-ray response. Together, these results indicate that the fixed clinical RBE of 1.1 may reasonably approximate some proton irradiation conditions but does not fully capture the range of biological responses observed across different beam positions, irradiation fields, cell lines, and surviving fraction endpoints. These observations support recent task group recommendations that LET_D_ should be considered when interpreting biological effect,^13^ while also emphasizing that LET_D_ alone does not fully characterize biological response across all irradiation conditions.

Carbon ion irradiation produced a substantially greater increase in RBE than proton irradiation, with RBE_10_ increasing markedly across the sampled pristine beam LET_D_ range. In PANC-1, RBE_10_ rose from approximately 2.3 in the entrance region to 5.2 at the Bragg peak, while in BxPC-3 it increased from 1.3 to 3.7 across the same range. These values are broadly consistent with previous reports of increasing carbon ion RBE with LET_D_ in pancreatic cancer models. Matsui et al. reported RBE values ranging from approximately 1.2–1.8 at 13 keV/µm to 1.8–2.5 at 80 keV/µm across MIAPaCa-2, SUIT-2, and BxPC-3 cell lines.^17^ Similarly, raster-scanned carbon ion experiments at HIT demonstrated RBE values of 1.5 – 4.5 for AsPC-1, BxPC-3 and PANC-1 cells irradiated within an extended Bragg peak with an LET_D_ of 103 keV/µm.^19^ Oonishi et al. likewise reported RBE values near 2 for MIA PaCa-2 and BxPC-3 exposed to an SOBP with LET values around 50 keV/µm.^11^ Collectively, these studies and the present findings support an association between increasing LET_D_ and greater carbon ion biological effectiveness, while also demonstrating substantial variation with cell model, irradiation conditions, and RBE endpoint.

An important distinction emerged when comparing pristine and SOBP carbon ion irradiations. Although the carbon ion SOBP and a proximal pristine position had similar LET_D_ values, the SOBP produced lower RBE values in both cell lines. Comparable LET_D_ therefore did not translate into equivalent biological effectiveness across the two irradiation fields. This difference was not captured by LET_D_ alone and may be related to differences in particle and energy spectra, fragment contributions, or microdosimetric distributions between the irradiation fields, consistent with previous theoretical findings.^27-29^ Because the present study was not designed to isolate the contribution of these individual physical factors, their relative roles cannot be determined from these data. Nevertheless, the findings indicate that LET_D_ alone may not fully characterize biological effectiveness across different field compositions.

### Changes in linear-quadratic parameters with increasing dose-averaged linear energy transfer

The reduced curvature of the survival curves at higher LET_D_ reflects the increasingly linear dose-response characteristic of densely ionizing radiation. This response is consistent with the greater probability of lethal damage from individual particle tracks at higher LET.^30^ These changes in survival curve shape also produced surviving fraction-dependent variation in RBE, with the reduced shoulder relative to the X-ray reference resulting in higher RBE values at higher surviving fractions.

### Cell-line-specific differences in carbon ion response

The clearest difference between PANC-1 and BxPC-3 was evident at the highest LET_D_ pristine carbon ion position. At matched absorbed doses, PANC-1 showed lower clonogenic survival than BxPC-3, with an approximately tenfold difference at 3 Gy. The corresponding carbon ion RBE values were also higher in PANC-1, indicating that increasing LET_D_ did not produce equivalent biological response in the two cell lines. Because both cell lines were irradiated using the same geometry and dosimetry, the divergence suggests that intrinsic cellular characteristics contributed to the different responses observed at high LET_D_. In contrast, the carbon ion SOBP RBE values were similar between the cell lines. Pristine beam measurements may therefore reveal cell-line-specific differences that are less apparent following exposure to the mixed particle and energy spectra of an SOBP, although further investigation is necessary to determine which physical components of the field contribute to this difference.

PANC-1 and BxPC-3 differ in several biological characteristics that may provide context for their responses.^19^ PANC-1 harbors KRAS^G12D^ and TP53 mutations and exhibits mesenchymal-like characteristics, whereas BxPC-3 retains epithelial-like features and is KRAS wild-type.^31^ Differences in DNA damage response, repair–related signaling including pathways involving 14-3-3σ, apoptotic regulation, and cell death pathway activation have also been reported between these cell lines.^18, 32-38^ These characteristics provide plausible biological context for the different carbon ion responses but were not evaluated mechanistically in the present study.

### Implications for relative biological effectiveness modeling and dose interpretation

The dataset presented here provides survival curves and fitted α and β parameters across pristine proton and carbon ion beam positions, together with measurements under mid-SOBP irradiation. These data provide an experimental reference for future evaluation of RBE models across the physical and biological conditions represented here. The pristine beam measurements span a broad LET_D_ range, allowing comparisons of predicted and observed RBE, changes in α and survival-curve curvature at high LET_D_, and differences between the two cell lines. The SOBP measurements extend this comparison to mixed particle and energy spectra.

The alternative LET_D_ definitions and microdosimetric quantities reported here allow evaluation of whether information beyond a single LET_D_ value improves characterization of biological response across pristine and SOBP irradiation. Similarly, reproducing the increase in RBE with LET_D_ is not sufficient if a model does not also capture the differing response between cell lines. Together, these measurements provide reference data for determining where current biological and physical assumptions reproduce the observed PDAC response and where additional inputs may be needed.

The clinical relevance of this work lies in the ongoing challenge of comparing carbon ion therapy outcomes for PDAC across institutions, where treatment delivery techniques and RBE models vary considerably.^2, 39^ A physically characterized dataset spanning both pristine and SOBP irradiation provides a common reference for evaluating how these methodological differences influence estimates of biological effect. This is particularly important in PDAC, where regions of elevated LET may occur within the target tumor but can also extend into adjacent radiosensitive gastrointestinal tissues. Although these *in vitro* findings cannot be translated directly into predictions of tumor control or normal-tissue toxicity, they provide important radiobiological context for interpreting and comparing emerging clinical outcomes.

### Limitations and future work

This study evaluated two PDAC cell lines in two-dimensional monolayer culture and therefore does not represent the full biological and microenvironmental heterogeneity of pancreatic tumors. Clonogenic survival was the sole endpoint; no mechanistic assays were performed to determine whether differences in response were related to DNA damage complexity, repair capacity, apoptosis, or other cell-specific processes. The molecular features discussed above should therefore be considered biological context rather than demonstrated mechanisms.

The irradiation conditions also sampled four discrete positions along each pristine Bragg curve and a single central position within each SOBP. These measurements do not provide a continuous biological profile across the beam or characterize variation throughout the full modulated field. Although microdosimetric quantities were calculated, the study was not designed to measure or isolate the contribution of each physical component to the measured biological response.

Future work should extend these measurements to a broader panel of PDAC models, incorporate three-dimensional and microenvironmentally relevant systems,^40^ and include biological endpoints that can directly evaluate radiation damage and repair. Applying established and emerging RBE models to the present dataset will also be important for determining which physical and biological inputs are needed to reproduce and predict responses across pristine and SOBP irradiation, supporting the development of biologically informed ion therapy planning approaches.

This study establishes a physically characterized dataset linking clonogenic response and RBE to radiation quality in two PDAC cell models irradiated with pristine proton and carbon ion beams, complemented by measurements under SOBP irradiation conditions. Proton RBE remained near 1.1 across most sampled conditions but increased at the distal edge, whereas carbon ion RBE showed greater dependence on radiation quality and differed between PANC-1 and BxPC-3. Comparable dose-averaged LET_D_ values did not necessarily yield equivalent biological responses in pristine and SOBP carbon ion fields. These findings indicate that RBE in PDAC is influenced not only by LET_D_, but also by irradiation-field composition and cell-specific factors, and they provide experimentally grounded benchmark data for the evaluation and refinement of biological-effect models across distinct proton and carbon ion irradiation conditions.

## FUNDING

This work was supported by the State of Florida Casey DeSantis funds to Mayo Clinic in Florida. The authors gratefully acknowledge support from the Austrian Federal Ministry of Women, Science and Research (BMBWF) for beam time at MedAustron.

## ACKNOWLEDGEMENTS

ChatGPT (GPT-5.6, OpenAI) was used for final editorial review. All edits were reviewed by the authors; the scientific content, phrasing, and interpretation are the authors’ own.

## CONFLICT OF INTEREST

The authors declare no conflicts of interest.

## Notes

### Competing Interest Statement

The authors have declared no competing interest.

