## Supplementary Materials for "Clonogenic survival and relative biological effectiveness of pancreatic ductal adenocarcinoma under pristine and spread-out Bragg peak irradiation with protons and carbon ions"

*Seeding Densities*

Supplementary Figure 1 summarizes the dose- and cell-line-dependent seeding densities used for clonogenic survival analysis following X-ray, pristine proton, pristine carbon-ion, and SOBP irradiation. Seeding densities were selected to maintain countable colony numbers (aiming for 80-120 cells per well) across the expected range of surviving fractions. Sham-handled and 0-Gy samples were seeded at the corresponding control density and were used to determine plating efficiency for survival normalization.

| Cell Line | Radiation | Position | Seeding Density | | | | | | |
|---|---|---|---|---|---|---|---|---|---|
| **PANC-1** | **X-ray** | | **0 Gy/SHAM** | **1 Gy** | **2 Gy** | **3 Gy** | **4 Gy** | **5 Gy** | **6 Gy** |
| | | | 500 | 500 | 500 | 2000 | 2000 | 2000 | 4000 |
| | **Proton** | | **0 Gy/SHAM** | **1 Gy** | **2 Gy** | **4 Gy** | **6 Gy** | **8 Gy** | |
| | | Entrance | 500 | 2000 | 2000 | 2000 | 4000 | 4000 | |
| | | Proximal | 500 | 2000 | 2000 | 2000 | 4000 | 4000 | |
| | | Peak | 500 | 2000 | 2000 | 2000 | 4000 | 4000 | |
| | | Distal | 500 | 2000 | 2000 | 2000 | 4000 | 4000 | |
| | | | **0 Gy/SHAM** | **0.5 Gy** | **1 Gy** | **2 Gy** | **4 Gy** | **6 Gy** | |
| | | Mid-SOBP | 500 | 500 | 2000 | 2000 | 2000 | 4000 | |
| | **Carbon** | | **0 Gy/SHAM** | **1 Gy** | **1.5 Gy** | **4 Gy** | **5 Gy** | **6.5 Gy** | |
| | | Entrance | 500 | 2000 | 2000 | 2000 | 4000 | 4000 | |
| | | | **0 Gy/SHAM** | **0.3 Gy** | **0.7 Gy** | **2 Gy** | **2.5 Gy** | **4 Gy** | |
| | | Proximal | 500 | 2000 | 2000 | 2000 | 4000 | 4000 | |
| | | | **0 Gy/SHAM** | **0.2 Gy** | **0.5 Gy** | **1.5 Gy** | **2 Gy** | **3 Gy** | |
| | | Pre-Peak | 500 | 2000 | 2000 | 2000 | 4000 | 4000 | |
| | | Peak | 500 | 2000 | 2000 | 2000 | 4000 | 4000 | |
| | | | **0 Gy/SHAM** | **0.5 Gy** | **1 Gy** | **2 Gy** | **4 Gy** | **6 Gy** | |
| | | Mid-SOBP | 500 | 500 | 2000 | 2000 | 2000 | 4000 | |
| **BxPC-3** | **X-ray** | | **0 Gy/SHAM** | **1 Gy** | **2 Gy** | **4 Gy** | **6 Gy** | **8 Gy** | |
| | | | 1250 | 2500 | 2500 | 5000 | 20000 | 20000 | |
| | **Proton** | | **0 Gy/SHAM** | **1 Gy** | **2 Gy** | **4 Gy** | **6 Gy** | **8 Gy** | |
| | | Entrance | 1250 | 2500 | 2500 | 5000 | 20000 | 20000 | |
| | | Peak | 1250 | 2500 | 2500 | 5000 | 20000 | 20000 | |
| | | Distal | 1250 | 2500 | 2500 | 5000 | 20000 | 20000 | |
| | | | **0 Gy/SHAM** | **0.5 Gy** | **1 Gy** | **2 Gy** | **4 Gy** | **6 Gy** | |
| | | Mid-SOBP | 500 | 2500 | 2500 | 5000 | 10000 | 10000 | |
| | **Carbon** | | **0 Gy/SHAM** | **1 Gy** | **1.5 Gy** | **4 Gy** | **5 Gy** | **6.5 Gy** | |
| | | Entrance | 1250 | 2500 | 2500 | 5000 | 20000 | 20000 | |
| | | | **0 Gy/SHAM** | **0.3 Gy** | **0.7 Gy** | **2 Gy** | **2.5 Gy** | **4 Gy** | |
| | | Proximal | 1250 | 2500 | 2500 | 5000 | 20000 | 20000 | |
| | | | **0 Gy/SHAM** | **0.2 Gy** | **0.5 Gy** | **1.5 Gy** | **2 Gy** | **3 Gy** | |
| | | Pre-Peak | 1250 | 2500 | 2500 | 5000 | 20000 | 20000 | |
| | | Peak | 1250 | 2500 | 2500 | 5000 | 20000 | 20000 | |
| | | | **0 Gy/SHAM** | **0.5 Gy** | **1 Gy** | **2 Gy** | **4 Gy** | **6 Gy** | |
| | | Mid-SOBP | 2500 | 2500 | 2500 | 2500 | 5000 | 20000 | |

**Supplementary Figure 1. Clonogenic seeding densities for PANC-1 and BxPC-3 cells by radiation quality and dose group**. Number of cells seeded per well for clonogenic survival assays under photon (X-ray), 180.1 MeV proton, 352.4 MeV/u carbon ion, and proton and carbon ion SOBP irradiation conditions. Seeding densities were adjusted according to dose level to maintain countable colony formation across decreasing surviving fractions. Sham and 0 Gy controls were included for each radiation condition to determine plating efficiency for normalization of survival curves.

*Bragg peak localization and sampling-position selection*

The water-equivalent thickness of the phantom entrance window (0.355 mm) and the geometric offset associated with the physical chamber positioning in the water tank (19.2 mm) were incorporated into the determination of measurement depths. A locking bar was indexed on the treatment couch to ensure reproducible phantom placement across measurement campaigns. All chamber readings were corrected for temperature, pressure, and beam quality for proton and carbon ion measurements.

An 8 Gy pre-irradiation was delivered prior to peak mapping to stabilize the electrometer and cabling. Depth-dose measurements were first acquired in coarse steps around the expected Bragg peak location and then refined in progressively smaller increments to determine the precise peak position. The Bragg peak was mapped using the Advanced Markus chamber. Monte Carlo simulations of the delivered beam were used to determine the relationship between $LET_D$ and distance from the Bragg peak. Combining the measured Bragg peak position with the simulated $LET_D$-depth relationship allowed selection of measurement depths corresponding to the targeted radiation-quality regions. This approach accounted for small differences between measured and simulated Bragg peak positions. Measurement depths were selected to achieve high $LET_D$ while limiting positional uncertainty, because the steeper $LET_D$ gradients near the distal region increase sensitivity to positioning uncertainty. The Bragg peak was mapped prior to each irradiation session to confirm positioning.

*Monte Carlo dose and $LET_D$ calculations*

The physical models used in OpenTOPAS differed between proton and carbon ion beams. For proton simulations, the following modules were used: g4em-standard_opt4, g4h-phy_QGSP_BIC_HP, g4decay, g4ion-binarycascade, g4h-elastic_HP, and g4stopping. For carbon ion simulations, the corresponding modules were g4em-standard_opt4, g4h-phy_QGSP_BIC_HP, g4decay, g4ion-liqmd, g4h-elastic_HP, and g4stopping. Following a methodology based on the reciprocity theorem, depth-dependent absorbed dose, dose-averaged LET ($LET_D$), and lineal-energy distributions at the center of 200 × 200 $mm^2$ radiation fields were estimated by laterally integrating pencil-beam results over 60 × 60 × 0.1 $mm^3$ computational phantom slices. Each beam simulation used $10^7$ primary ions. Absorbed dose was scored using the DoseToMedium scorer, and a validated in-house scorer was used to estimate mid-step $LET_D$ for user-selected ion types.

$RBE_{50}$, $RBE_{10}$, and $RBE_1$ were additionally examined using the alternative $LET_D$ definitions and dose-mean lineal energy ($\bar{y}_{D,all}$) (Supplementary Figures 3–7) to evaluate the influence of radiation-quality descriptor selection on the apparent relationship between radiation quality and RBE.

*Microdosimetric calculations*

Lineal-energy distributions (y) were scored using a revised implementation of a published TOPAS extension[1] based on an analytical microdosimetric function (AMF).[2] The AMF parameterizes detailed track-structure calculations performed with PHITS and computes microdosimetric distributions as a function of ion charge, kinetic energy, and LET. Unlike the original implementation, which relied on a pre-step LET lookup table, the LET used for the AMF calculations was evaluated at mid-step using ComputeElectronicDEDX. The resulting lineal-energy distributions included contributions from secondary electrons set in motion by the ions; secondary-electron transport was therefore disabled to avoid double counting. A target radius of 0.3 μm was used, consistent with previous studies.[3] Each simulation was parallelized across 256 processor threads on two Intel Xeon Platinum 8592+ CPUs; calculation times were approximately 13 h for the proton beam and 33 h for the carbon ion beam.

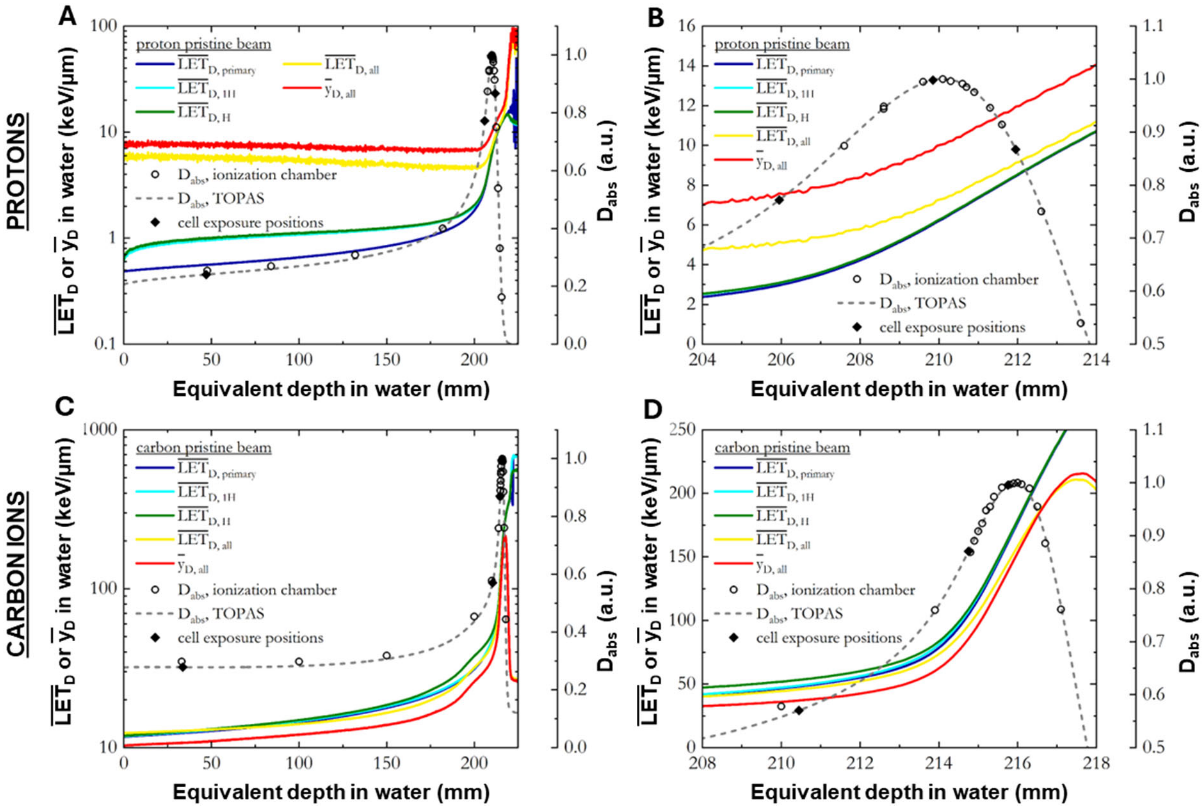


**Supplementary Figure *2*. Depth–dose and LET profiles for monoenergetic proton and carbon beams with selected biological sampling positions.** (A–B) Pristine depth–dose and LET distributions for the 180.1 MeV proton beam shown across the full range (A) and magnified near the Bragg peak region (B). (C–D) Corresponding distributions for the 352.4 MeV/u carbon ion beam shown across the full range (C) and magnified near the Bragg peak (D). Open circles represent measured depth–dose values acquired using the Advanced Markus ionization chamber; dashed curves indicate Monte Carlo (TOPAS) simulations. Solid lines show dose-averaged LET ($LET_D$) calculated under different ion-inclusion conditions (primary only, hydrogen/carbon fragments, and all ions), and red curves denote dose-averaged lineal energy ($\bar{y}_D$). Black diamonds indicate the four cell exposure depths selected per beam.

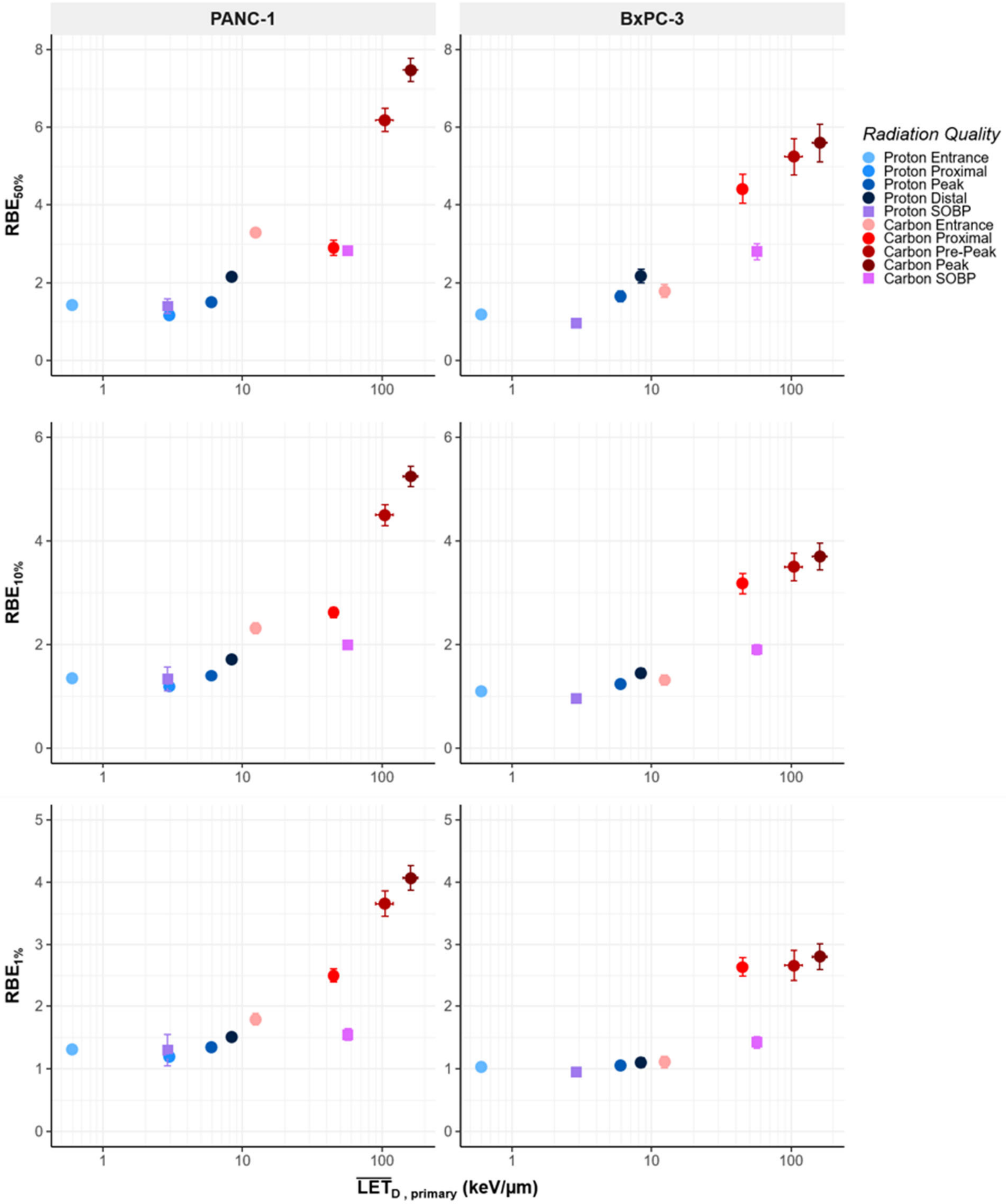


***Supplementary Figure 3. RBE$_{50}$, RBE$_{10}$, and RBE$_{1}$ as a function of LET$_{D,primary}$.*** Relative biological effectiveness calculated at 50%, 10%, and 1% surviving fraction (top, middle, and bottom rows, respectively) plotted against Monte Carlo–derived dose-averaged LET of primary ions only (LET$_{D,primary}$). Left panels show PANC-1 and right panels show BxPC-3 cells. Blue symbols represent 180.1 MeV proton sampling positions (entrance, proximal, peak, distal), and red symbols represent 352.4 MeV/u carbon ion sampling positions (entrance, proximal, pre-peak, peak). Purple and pink squares represent SOBP proton and carbon results, respectively. Error bars reflect propagated uncertainty from the fitted LQ parameter estimates used in the RBE calculation.

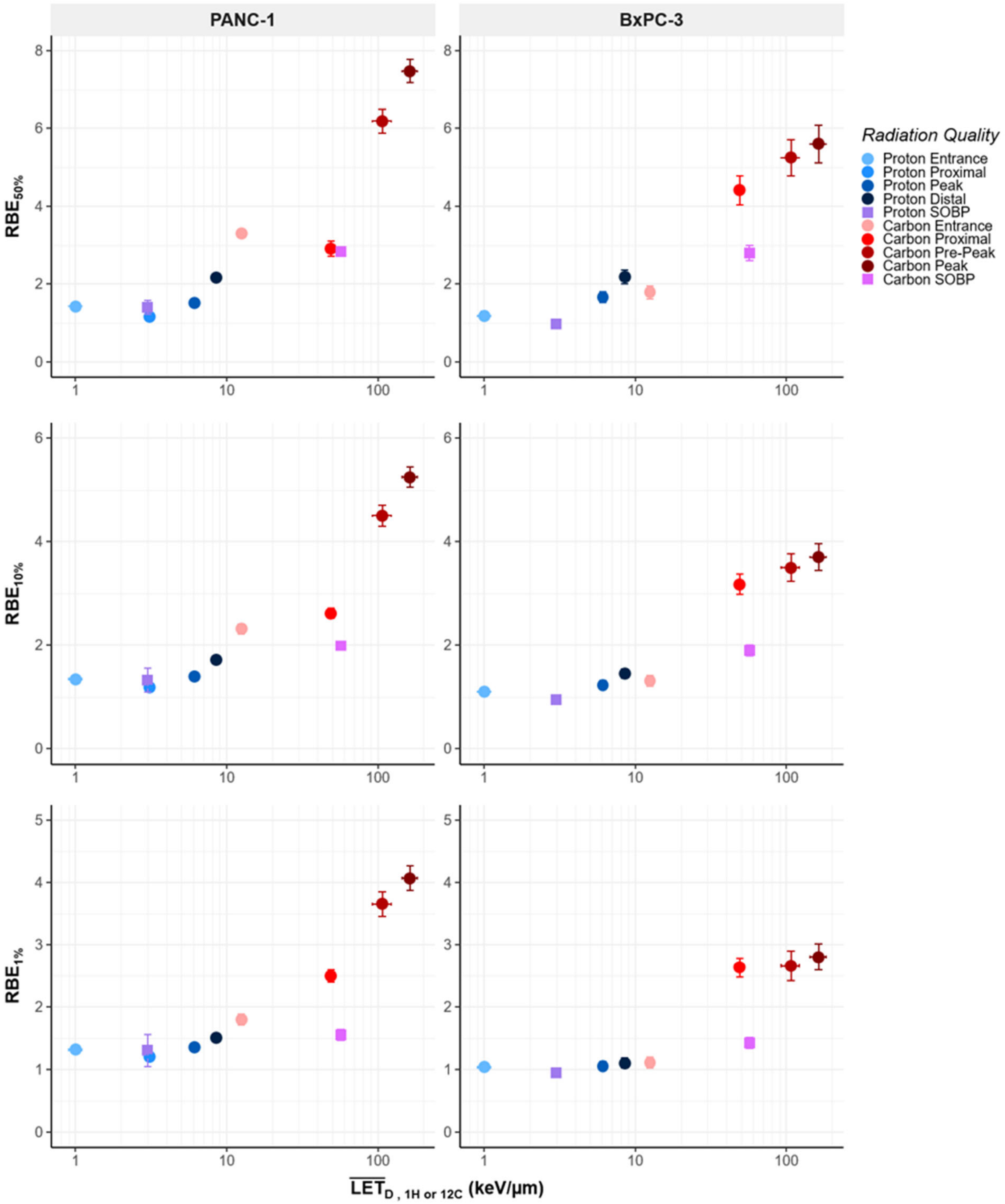


***Supplementary Figure 4. RBE₅₀, RBE₁₀, and RBE₁ as a function of LET$_{D,\,^{1}H\ or\ ^{12}C}$.*** Relative biological effectiveness at 50%, 10%, and 1% survival (top to bottom rows) plotted against dose-averaged LET including primary ions and same-ion fragments (LET$_{D,^{1}H}$ for protons or LET$_{D,^{12}C}$ for carbon ions). Left panels show PANC-1 and right panels show BxPC-3 cells. Symbols and color coding correspond to radiation quality and sampling position as indicated in legend. Error bars reflect propagated uncertainty from the fitted LQ parameter estimates used in the RBE calculation.

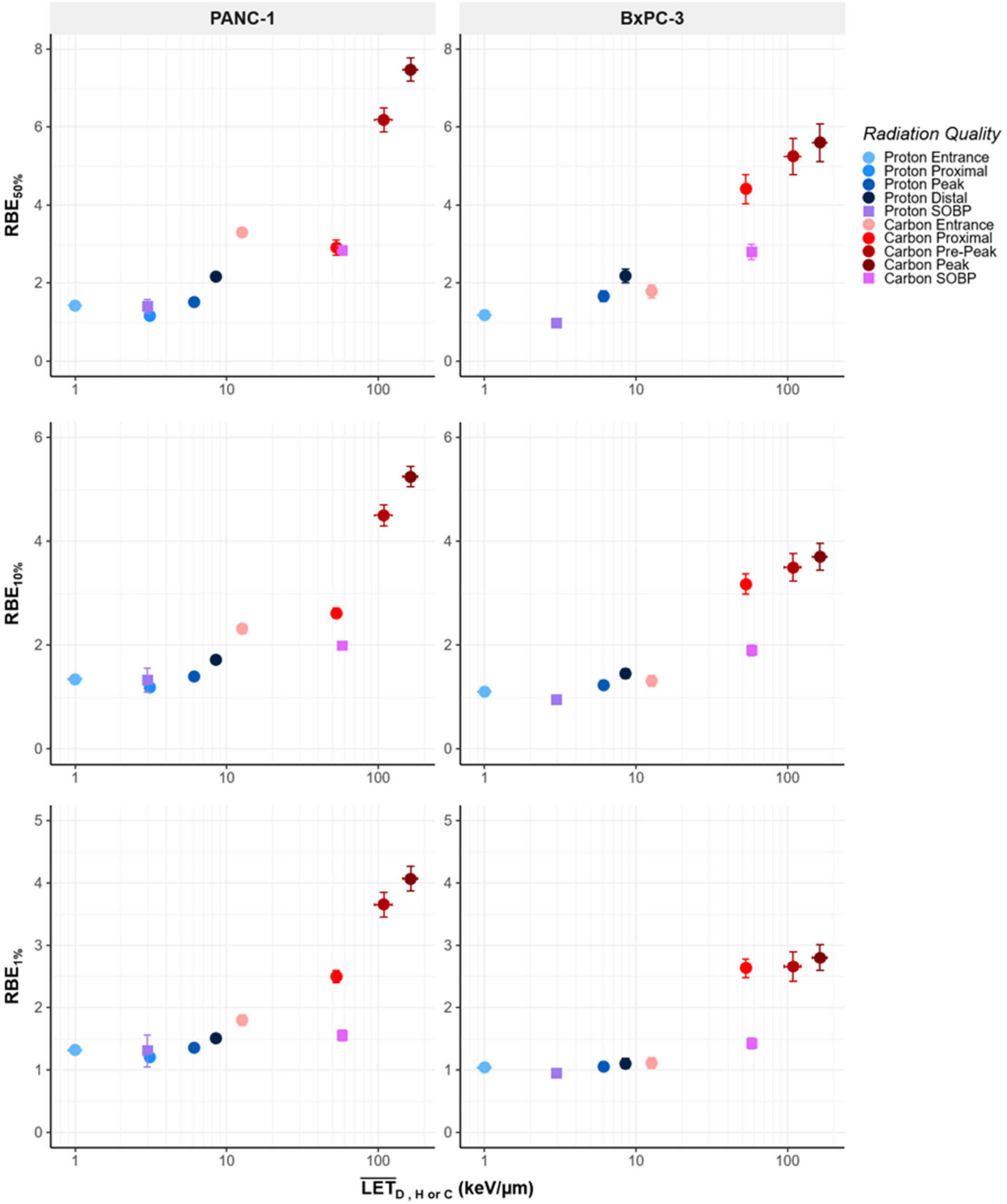


***Supplementary Figure 5. RBE$_{50}$, RBE$_{10}$, and RBE$_{1}$ as a function of LET$_{D,H\ or\ C}$.*** Relative biological effectiveness at 50%, 10%, and 1% survival (top to bottom rows) plotted against dose-averaged LET including primary ion and same-element fragments (LET$_{D,H}$ ) for proton irradiations and (LET$_{D,C}$) for carbon irradiations. Left panels show PANC-1 and right panels show BxPC-3 cells. Symbols and color coding correspond to radiation quality and sampling position as indicated in legend. Error bars reflect propagated uncertainty from the fitted LQ parameter estimates used in the RBE calculation.

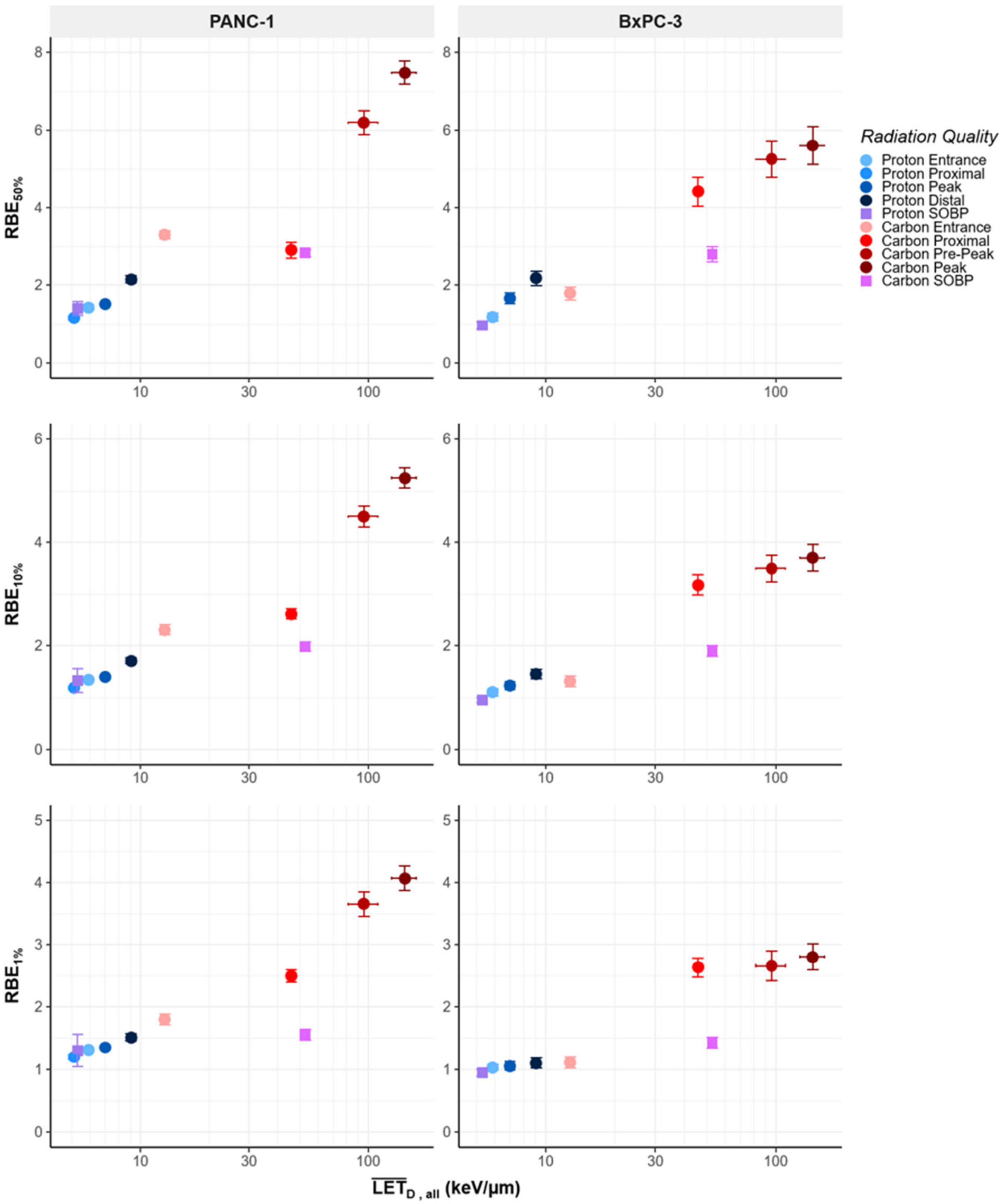


***Supplementary Figure 6. $RBE_{50}$, $RBE_{10}$, and $RBE_1$ as a function of $LET_{D,all}$.*** Relative biological effectiveness at 50%, 10%, and 1% survival (top to bottom rows) plotted against dose-averaged LET including all ion contributions ($LET_{D,all}$). Left panels show PANC-1 and right panels show BxPC-3 cells. Symbols and color coding correspond to radiation quality and sampling position as indicated in legend. Error bars reflect propagated uncertainty from the fitted LQ parameter estimates used in the RBE calculation.

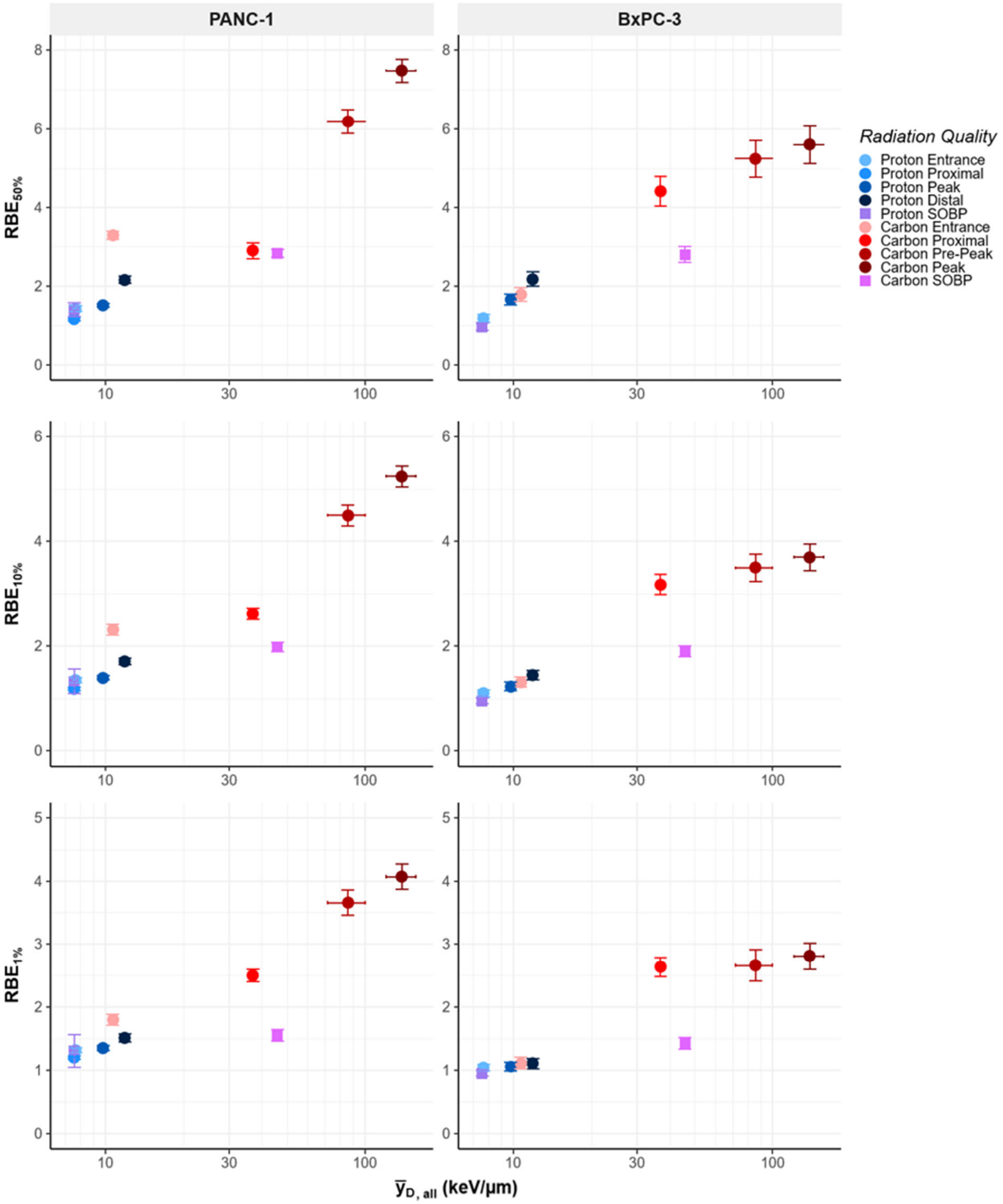


***Supplementary Figure 7. RBE$_{50}$, RBE$_{10}$, and RBE$_{1}$ as a function of dose-averaged lineal energy ($\bar{y}_{D,all}$).*** Relative biological effectiveness at 50%, 10%, and 1% survival (top to bottom rows) plotted against Monte Carlo–derived dose-averaged lineal energy ($\bar{y}_D$). Left panels correspond to PANC-1 and right panels correspond to BxPC-3 cells. Symbols and color coding correspond to radiation quality and sampling position as indicated in legend. Error bars reflect propagated uncertainty from the fitted LQ parameter estimates used in the RBE calculation.
